# Acute stress impairs sensorimotor gating via the neurosteroid allopregnanolone in the prefrontal cortex

**DOI:** 10.1101/2022.06.05.494755

**Authors:** Roberto Cadeddu, Laura J Mosher, Peter Nordkild, Nilesh Gaikwad, Gian Michele Ratto, Simona Scheggi, Marco Bortolato

## Abstract

**Background and purpose:** Ample evidence indicates that environmental stress impairs information processing, yet the underlying mechanisms remain partially elusive. We showed that, in several rodent models of psychopathology, the neurosteroid allopregnanolone (AP) reduces the prepulse inhibition (PPI) of the startle, a well-validated index of sensorimotor gating. Since this GABA_A_ receptor activator is synthesized in response to acute stress, we hypothesized its participation in stress-induced PPI deficits.

**Experimental approach:** We studied whether and how AP influences PPI in mice and rats; thereafter, we tested AP’s implication in the PPI deficits produced by several complementary regimens of acute and short-term stress (footshock, restraint, predator exposure, and sleep deprivation).

**Key results:** Systemic AP administration reduced PPI in C57BL/6J mice and Long-Evans, but not Sprague-Dawley, rats. These effects were reversed by isoallopregnanolone (isoAP), an endogenous AP antagonist, and the GABA_A_ receptor antagonist bicuculline and mimicked by AP infusions in the medial prefrontal cortex (mPFC). PPI was reduced by acute footshock, sleep deprivation as well as the combination of restraint and predator exposure in a time- and intensity-dependent fashion. Acute stress increased AP concentrations in the mPFC, and its detrimental effects on PPI were countered by systemic and intra-mPFC administration of isoAP.

**Conclusions and implications:** These results collectively indicate that acute stress impairs PPI by increasing AP content in the mPFC. The confirmation of these mechanisms across distinct animal models and several acute stressors strongly supports the translational value of these findings and warrants future research on the role of AP in information processing.

## INTRODUCTION

Sensorimotor gating is the perceptual function aimed at filtering out irrelevant and redundant contextual information. This selection process plays a crucial role in enabling the formation of saliency maps and enacting adaptive behavioral responses to environmental cues. The best-validated index to measure the integrity of sensorimotor gating is the prepulse inhibition (PPI) of the startle reflex (Swerdlow et al., 1992), defined as the reduction of the startle response that occurs when the eliciting acoustic pulse is immediately preceded by a weaker prestimulus (Graham et al., 1975). PPI deficits are a transdiagnostic feature of several psychopathological conditions (Geyer, 2006), including schizophrenia (Braff et al., 1978), mania (Perry et al., 2001), and Tourette syndrome (TS) (Castellanos et al., 1996). Notably, PPI is a cross-species endophenotype, which is also reduced in animal models of these neuropsychiatric disorders (Powell et al., 2009; Young et al., 2010; Godar et al., 2014). Several psychotomimetic drugs have been shown to damage PPI, including dopaminergic agonists and NMDA glutamate receptor antagonists (Geyer et al., 2001). Notably, the effects of some of these drugs are reversed by antipsychotics (Geyer et al., 2001), even though different species and strains exhibit differential sensitivities to the activation of D_1_ and D_2_ dopamine receptors (Mosher et al., 2016).

In addition to drug-based interventions, PPI can be reduced by stressful manipulations. Studies on animal models have pointed to the detrimental effects of some, but not all, stressors on PPI. For example, studies in Sprague-Dawley rats have shown that PPI is potently reduced by sleep deprivation (Frau et al., 2008) and exposure to a predator, but not by footshock (Bakshi et al., 2011). In other rat strains, chronic but not acute restraint stress was found to reduce PPI (Sutherland and Conti, 2011). Understanding the impact of stress on gating functions is particularly important, given that impaired stress coping is a common feature of neuropsychiatric disorders featuring gating deficits (Hoehn-Saric et al., 1995; Myin-Germeys et al., 2003; Myin-Germeys and van Os, 2007; Lataster et al., 2009; Conelea et al., 2011; Cougle et al., 2013). Although preliminary studies indicate that corticotropin-releasing factor (Conti et al., 2002; Bakshi et al., 2011) plays a role in these deficits, the underlying mechanisms remain poorly understood.

Over the past few years, our group has shown that PPI is modulated by neurosteroids, a family of brain-borne steroids (Baulieu, 1981) that play a critical role in the modulation of the stress response (Purdy et al., 1991; Girdler et al., 2012; Gunn et al., 2015). For example, our studies have shown that the modulation of neurosteroidogenic enzyme 5α-reductase modulates the response to dopaminergic agonists in rats and mice (Bortolato et al., 2008; Devoto et al., 2012; Frau et al., 2013; Frau et al., 2016). We recently documented that these imbalances are contributed by the neurosteroid allopregnanolone (AP; 3α,5α-tetrahydroprogesterone) (Mosher et al., 2019), a progesterone metabolite acting as a potent positive allosteric modulator of GABA_A_ receptors (Majewska et al., 1986; Puia et al., 1990). In animal models, brain AP synthesis is increased by several acute and short-term stressors (Purdy et al., 1991; Vallee et al., 2000), including sleep deprivation (Frau et al., 2017), restraint (Higashi et al., 2005) and footshock (Serra et al., 2002).

Building on these premises, we hypothesized that AP might mediate the PPI-disrupting effects of acute stress. To test this possibility, here we tested whether AP may disrupt PPI in rodents and through what mechanisms; furthermore, we examined whether blocking AP signaling may oppose the sensorimotor gating impairments induced by acute or short-term stressors.

## MATERIALS AND METHODS

### Animals

Experiments were performed using adult male mice (3-4-month-old) and rats (2-3-month-old). C57BL/6J mice (RRID: IMSR_JAX:000664) were purchased from The Jackson Laboratory (Bar Harbor, ME, USA). Long-Evans (RRID: RGD_2308852) and Sprague-Dawley rats (RRID: MGI:5651135) were obtained from Charles River Laboratories (Hollister, CA, USA). All animals were allowed to acclimate to facilities for 7-10 days before starting experimental procedures. Housing facilities were maintained at 22 °C with a 12-hour light/dark cycle (lights on at 6:00 AM hour, lights off at 6:00 PM hour). Experimental manipulations were carried out during the light cycle (between 9:00 AM and 5:00 PM). In conformity with the 3Rs principles (Russell and Burch, 1959), every effort was made to minimize the number and suffering of animals. Thus, for each experiment, animal numbers were determined based on power analyses conducted on preliminary results. All handling and manipulation procedures were performed in compliance with the National Institute of Health and ARRIVE guidelines (Percie du Sert et al., 2020) and approved by the local Institutional Animal Care and Use Committees. Animal studies are reported in compliance with the recommendations made by the British Journal of Pharmacology (Lilley et al., 2020). Studies were only limited to male animals to control the significant variations of AP levels in relation to the menstrual cycle (Schmidt et al., 1994; Genazzani et al., 1995; Palumbo et al., 1995)

### Drugs

AP was purchased from Tocris Bio-Techne (Minneapolis, MN, USA). For systemic injections, AP was dissolved in 2.5% DMSO, 2.5% Tween 80, and 0.9% NaCl. For intracerebral infusions, AP was dissolved in Tween 80/Ringer solution (final concentration, 1:1, v:v); AP was then infused in a solution of cyclodextrin/Ringer solution (final concentration 1:5, v:v). The 3β-epimer of AP, isoallopregnanolone (isoAP; 3β,5α, tetrahydroprogesterone) was donated by Asarina Pharma AB (Solna, Sweden) and suspended in 3% hydroxypropyl β-cyclodextrin for systemic injections. For intracerebral administrations, this suspension was further dissolved in Ringer solution (final concentration 1:1, v:v). SCH 23390, haloperidol, and bicuculline were obtained from Sigma-Aldrich (St. Louis, MO, USA). SCH 23390, haloperidol, and bicuculline were dissolved in 10% acetic acid buffered with NaOH and diluted with 0.9% NaCl. The injection volume for all systemic administrations was 10 ml/kg in mice and 2 ml/kg in rats.

### Stress procedures

The following acute and short-term stressors were used:

#### Foot shock

Foot shock stress was delivered through the Potentiated Startle Kit within sound-attenuating startle chambers (SR-LAB, San Diego Instruments, San Diego, CA, USA). Mice and rats were placed in Plexiglass enclosures with shock grids within sound-attenuating startle chambers. After a one-min acclimation period, animals were subjected to nine brief (1 s) electric shocks (0.2-0.5 mA), administered at random intervals of 30-60 s over 5 min.

#### Restraint

Mice were physically restrained for 30, 60, or 120 min inside a well-ventilated 50-ml plastic conical tube within a novel cage. Non-stressed control mice remained in an equivalent novel cage until PPI testing (Fig.1A). Rats were restrained in Plexiglass tubes (diameter: 7.5 cm; length: 30 cm; World Precision Instruments, Sarasota, FL) endowed with an adjustable end to fit the size of each animal.

**Figure 1.**
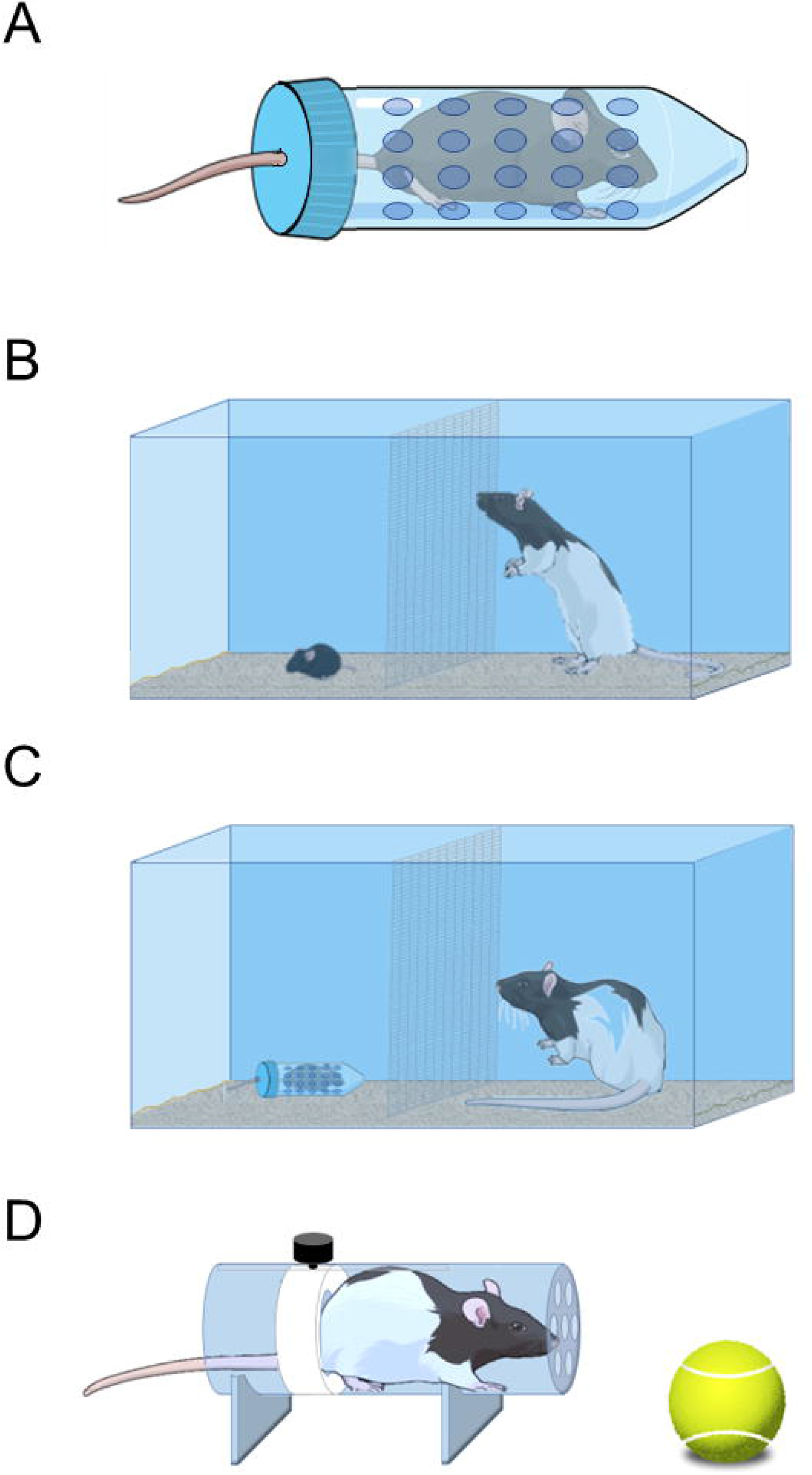
Schematic representation of some of the stressors used in the present study. A) Mice were restrained in plastic conical tubes with holes for ventilation. B) Mice were placed in one of the two compartments of a cage, with an awake rat on the opposite side. C) Restraint and predator stress was combined to increase the overall intensity of the stress imposed on mice. D) Rats were also subjected to a combination of restraint and predator cue exposure, using a standard cylindrical restrainer and a tennis ball previously impregnated with bobcat urine and placed in the vicinity of the rat’s snout. For further details, see text.

#### Predator exposure

Mice were placed in one of the two compartments of a novel cage (separated from each other by a metal grid; Fig. 1B). The opposite compartment hosted a LE male rat (weight: 225-250 g). This configuration allowed the mouse to see, smell, and hear the rat while avoiding direct physical contact (Fig. 1B). Exposure lasted 30, 60, or 120 min. The same setting was also used in combination with restraint stress to test the effects of the association of these two manipulations (Fig. 1C). The association of restraint and predator cue exposure test was also repeated in Long-Evans rats (Fig. 1D). To ensure predator cue exposure akin to that used for mice, restrained rats were placed near a tennis ball previously sprayed with 5 ml of bobcat urine (Fox Peak Outdoor Supply, Spring Grove, PA) (Fig. 1D) for 30 or 60 min.

#### Sleep deprivation

Sleep deprivation was obtained via the small-platform method, as previously described (Frau et al., 2008), a well-validated experimental paradigm that has been shown to produce near-complete REM sleep reduction and a marked reduction in overall sleep (Grahnstedt and Ursin, 1985). Mice and rats were kept on small Plexiglas platforms (4 and 7 cm in diameter, respectively) within a deep tank filled with water. Each platform was surrounded by water (up to 0.5 and 1 cm, respectively) beneath the surface. Food and water were freely available by placing chow pellets and water bottles on a grid located on the top of the tank. The temperature of the experimental room and the water inside the tank was maintained at 23±1°C. Control animals were placed in the experimental room but kept individually in their home cages after removing cage mates. Our prior studies showed that these animals exhibited the same behavioral phenotypes as those of rats kept on large platforms (Frau et al., 2008). SD duration was kept at 24 h for C57BL/6J mice and 48 h for Long-Evans rats, as these durations proved sufficient to produce stable PPI deficits. Both SD-subjected and NSD rats had free access to food and water throughout the procedure.

### PPI of the acoustic startle reflex

Startle testing was conducted in sound-attenuating startle chambers (SR-LAB, San Diego Instruments) as previously detailed (Bortolato et al., 2008). The startle testing protocol featured a 70 dB background white noise (5 minutes acclimatization period), followed by three consecutive blocks of pulse, prepulse + pulse, and “no stimulus” trials. During the first and third blocks, mice received five pulse-alone trials of 115 dB. During the second block, animals underwent a pseudorandom sequence of 50 trials, consisting of pulse-alone trials (twelve events), 30 trials of pulse preceded by 73, 76, 82 dB prepulse intensities (ten events for each level of prepulse loudness), and 8 no-stimulus trials. Intertrial intervals were selected randomly with a duration between 10 and 15 seconds. A dynamic calibration system was used to ensure comparable sensitivities across chambers. Sound levels were assessed using an A-scale setting. Percent PPI (%PPI) was calculated using the formula (Mean Startle Amplitude, MSA):

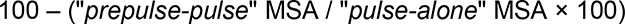

The first five “pulse-alone” bursts were excluded from the calculation (used as internal control). As no interactions between prepulse levels and treatment(s) were found in the statistical analysis, %PPI values were collapsed across prepulse intensity to represent average %PPI.

### Stereotaxic surgery and microinjections

Stereotaxic surgery was performed as previously described (Cadeddu et al., 2021). Briefly, mice were anesthetized with xylazine/ketamine (20/80 mg·kg^-1^, IP) and then placed onto a stereotaxic frame (David Kopf Instruments, Tujunga, CA, USA). Stainless steel 26-G bilateral guide cannulae (P1 Technologies Inc., Roanoke, VA, USA) were implanted in mPFC (AP + 1.8 mm, ML ± 0.3 mm, DV - 2.5 mm from the skull surface) or nucleus accumbens (NAc) (AP + 1.4 mm, ML ±0.5 mm, DV - 4.3 mm from the skull surface) according to the stereotaxic brain atlas (Paxinos and Watson, 1998). The lengths of the cannulae were selected to end 0.5 mm above the targeted area, and the injector projected 1 mm beyond the guide cannula. After implantation, cannulae were plugged with wire stylets to avoid clogging during the post-operative recovery period. Antibiotic therapy was administered for two days (enrofloxacin, Bayer HealthCare, Shawnee Mission, KS, USA). Mice were allowed to recover in their home cages (single housed) with food and water ad libitum. Four-five days after cannulation, mice received bilateral microinjections in the targeted area (according to the specific experimental design).

Microinjections were performed by gently restraining the mouse, removing the stylet, and replacing it with the injector (P1 Technologies Inc., Roanoke, VA, USA) connected by a 250 μl Hamilton syringes via PE tubing. A microinfusion pump (KD Scientific, Holliston, MA, USA) delivered 0.5 µl/side of drug solution (or its vehicle solution) at a constant flow/rate of 0.25 µl/min. After infusion, the injector was left in place for two minutes to allow fluid diffusion. PPI experiments were carried out immediately after infusion. After behavioral tests, animals were sacrificed, and the histological verification of cannula location was confirmed. Animals with errant locations of either device or damage to the targeted area(s) were excluded from the analysis.

### Neurosteroid quantification

Reference standards were purchased from Steraloids (Newport, RI, USA). Stock steroid standards were prepared by mixing 5 µl of 1 mg/ml solution of each steroid and adjusting the final volume to 1 ml using methanol. All the stock solutions were stored at −80 °C. Samples were extracted and purified as previously described (Gaikwad, 2013). Following extraction with 0.7 ml chloroform, samples were centrifuged, and the chloroform fraction was transferred to a 2-ml tube and dried. The resulting residue was again extracted with 0.5 ml methanol. The methanol fraction was then added to the above chloroform fraction; the mixture was dried and resuspended in 130 µl methanol, filtered, and filtrates were transferred to vials for ultra-performance liquid chromatography-tandem mass spectrometry (UPLC-MS/MS) analysis. UPLC-MS/MS analyses were performed using a Waters Acquity UPLC system connected with the high-performance triple quadrupole (TQD) mass spectrometer. Analytical separations were conducted as described in Bossé et al. (2021). The elution from the UPLC column was introduced to the TQD mass spectrometer. All MS experiments were performed by using electrospray ionization (ESI) in positive ion (PI) mode, with an ESI-MS capillary voltage of 4.0 kV, an extractor cone voltage of 3 V, and a detector voltage of 500 V. Desolvation gas was kept at 500 l/h, with temperature at 350 °C and source temperature at 150 °C. Pure standards of all targeted steroids were used to optimize the UPLC-MS/MS conditions before analysis and making calibration curves. Reference standards were run before the first sample, in the middle of the runs, and after the last sample to prevent errors due to matrix effect and day-to-day instrument variations. In addition, spiked samples were run before the first sample and after the last sample to calibrate for the drift in the retention time due to the matrix effect. After standard and spiked sample runs, several blanks were injected to wash the injector and avoid carry-over effects. The resulting data were processed using Waters Target Lynx 4.1 software (Bossé et al., 2021). The analytes concentrations were normalized per gram of tissue before extraction.

### Statistical analyses

Data and statistical analyses comply with the recommendations and requirements for experimental design and analysis outlined by Curtis et al. (2018). Normality and homoscedasticity of data distribution were verified using Kolmogorov-Smirnov and Bartlett’s tests. Statistical analyses of parametric data were performed using t-test, one-way ANOVA, or two-way ANOVA, as appropriate. *Post-hoc* comparisons were performed by Tukey’s test. Significance thresholds were set at 0.05.

## RESULTS

### AP dose-dependently reduces PPI in C57BL/6J mice and Long-Evans rats but not in Sprague-Dawley rats

We first tested the effects of systemic injections of AP (1, 3, 6, or 12 mg·kg^-1^, IP, 5 min before behavioral testing) in C57BL/6J mice. Although startle amplitude was not modified by any treatment (Fig. 2A), systemic administration of AP at 6 and 12 mg·kg^-1^ elicited a significant impairment of PPI (between-group, One-way ANOVA: F_4,43_=5.13, *p* < 0.05; *post-hoc* comparisons by Tukey’s; Fig. 2B). Similar to C57BL/6J mice, Long-Evans rats (Fig. 2C-D) treated with AP (12 mg·kg^-1^, IP) exhibited significant PPI impairments (unpaired t-test with Welch’s correction: t=2.880, df=9.819, *p* < 0.05; Fig. 2D) that were not paralleled by changes in startle modifications. In contrast with these findings, AP (12 mg·kg^-1^, IP) failed to modify startle amplitude and PPI in Sprague-Dawley rats (Fig. 2E-F).

**Figure 2.**
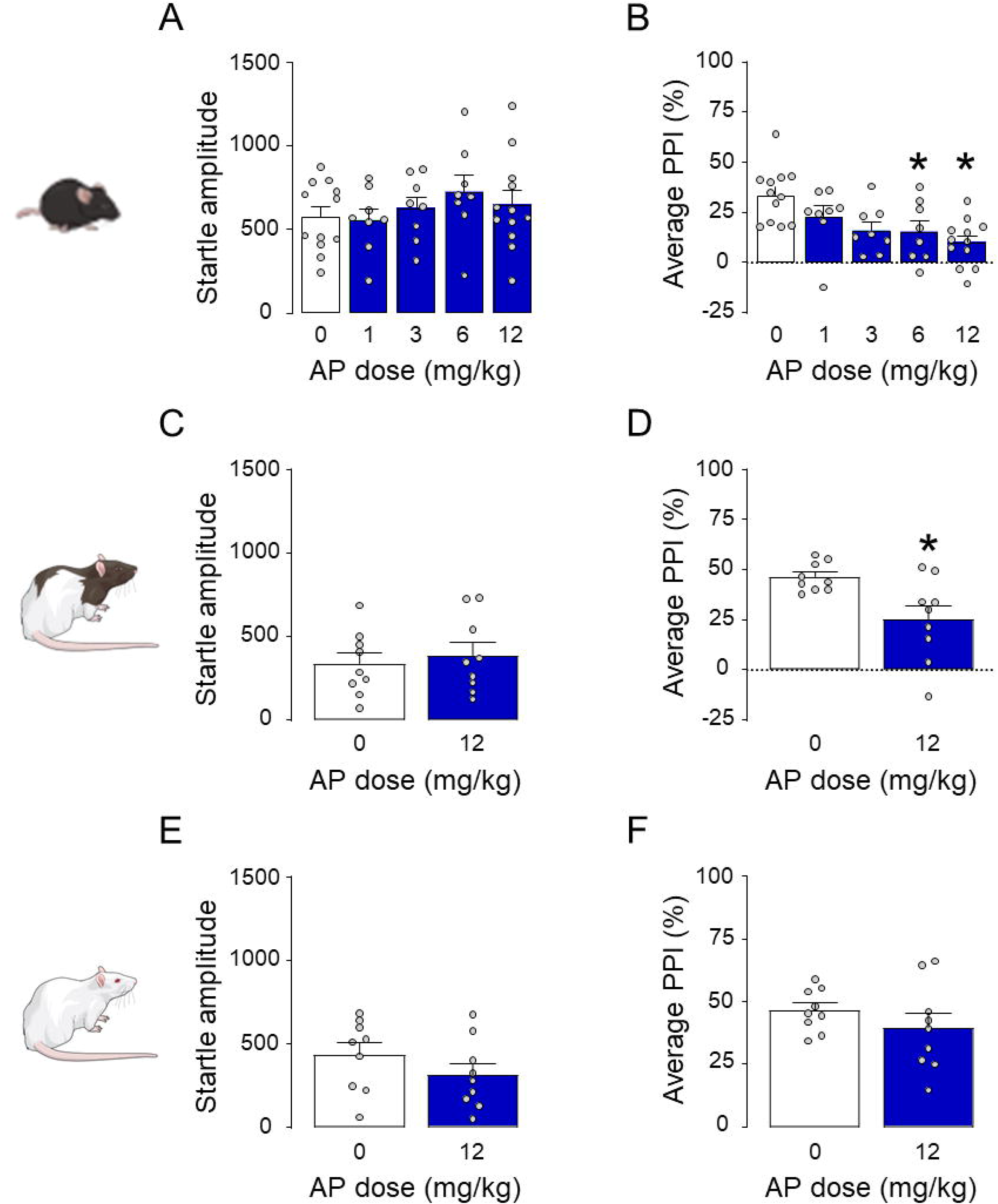
Effects of allopregnanolone (AP) administered systemically in C57BL/6J mice (A-B), Long-Evans rats (C-D), and Sprague-Dawley rats (E-F). AP doses are indicated in mg·kg^-1^, IP. Data are shown as means ± SEM; n=8-12/group. VEH, vehicle. *, *p* < 0.05 compared with 0 mg·kg^-1^ (VEH-treated). For further details, see text.

### The PPI-disrupting effects of AP are mediated by GABA_A_ but not dopamine receptors

We then tested whether the PPI disruption caused by AP was reversed by dopaminergic antagonists in C57BL/6J mice. As shown in Figs. 3A-B, AP (12 mg·kg^-1^, IP, 5 min before testing) significantly increased startle amplitude (Two-way ANOVA; Main AP effect: F_1,32_=7.24, *p* < 0.05) and reduced PPI (Two-way ANOVA; Main effect of AP: F_1,32_=8.56, *p* < 0.05), but neither effect was modified by the dopamine D_1_ receptor antagonist SCH23390 (0.5 mg·kg^-1^, SC, 5 min before AP). Similarly, the effects of AP were not opposed by the dopamine D_2_ receptor antagonist haloperidol (0.5 mg·kg^-1^, SC, 5 min before AP; Figs. 3C-D). Next, we studied whether the PPI deficits induced by AP were countered by systemic administration of the competitive GABA_A_ receptor antagonist bicuculline (1 mg·kg^-1^, IP, 5 min before AP; Figs. 3E-F). Bicuculline was able to oppose the detrimental effect of AP on sensorimotor gating (Two-way ANOVA; AP × bicuculline interaction: F_1,20_=7.39, *p* < 0.05; *p*’s <0.05 for comparisons of vehicle-vehicle vs. vehicle-AP and vehicle-AP vs. bicuculline-AP; Fig. 3F). Building on this evidence, we tested whether the PPI deficits induced by AP could be counteracted by isoAP (10 mg·kg^-1^, IP, 5 min before AP), the endogenous AP epimer acting as an antagonist of this steroid site on the GABA_A_ receptor. While AP increased the magnitude of the startle reflex (Two-way ANOVA; Main AP effect: F_1,43_=6.39, *p* < 0.05; Fig. 3G) and reduced PPI (Two-way ANOVA; Main AP effect: F_1,43_=22.41, *p* < 0.05; Fig. 3H), isoAP fully opposed the PPI impairment elicited by AP (Two-way ANOVA: AP × isoAP interaction: F_1,43_=34.24, *p* < 0.05; *p’*s < 0.05 for comparisons of vehicle-vehicle vs vehicle-AP and isoAP-AP vs vehicle-AP; Fig. 3H). In Long-Evans rats, isoAP (10 mg·kg^-1^, IP) induced no significant modification of the startle reflex (Fig. 4A) but fully countered the PPI deficits induced by systemic AP (12 mg·kg^-1^, IP) (Two-way ANOVA: AP × isoAP interaction: F_1,36_=6.42, *p* < 0.05; *p’*s < 0.05 for comparisons between vehicle-vehicle and vehicle-AP and between isoAP-AP and vehicle-AP; Fig. 4B).

**Figure 3.**
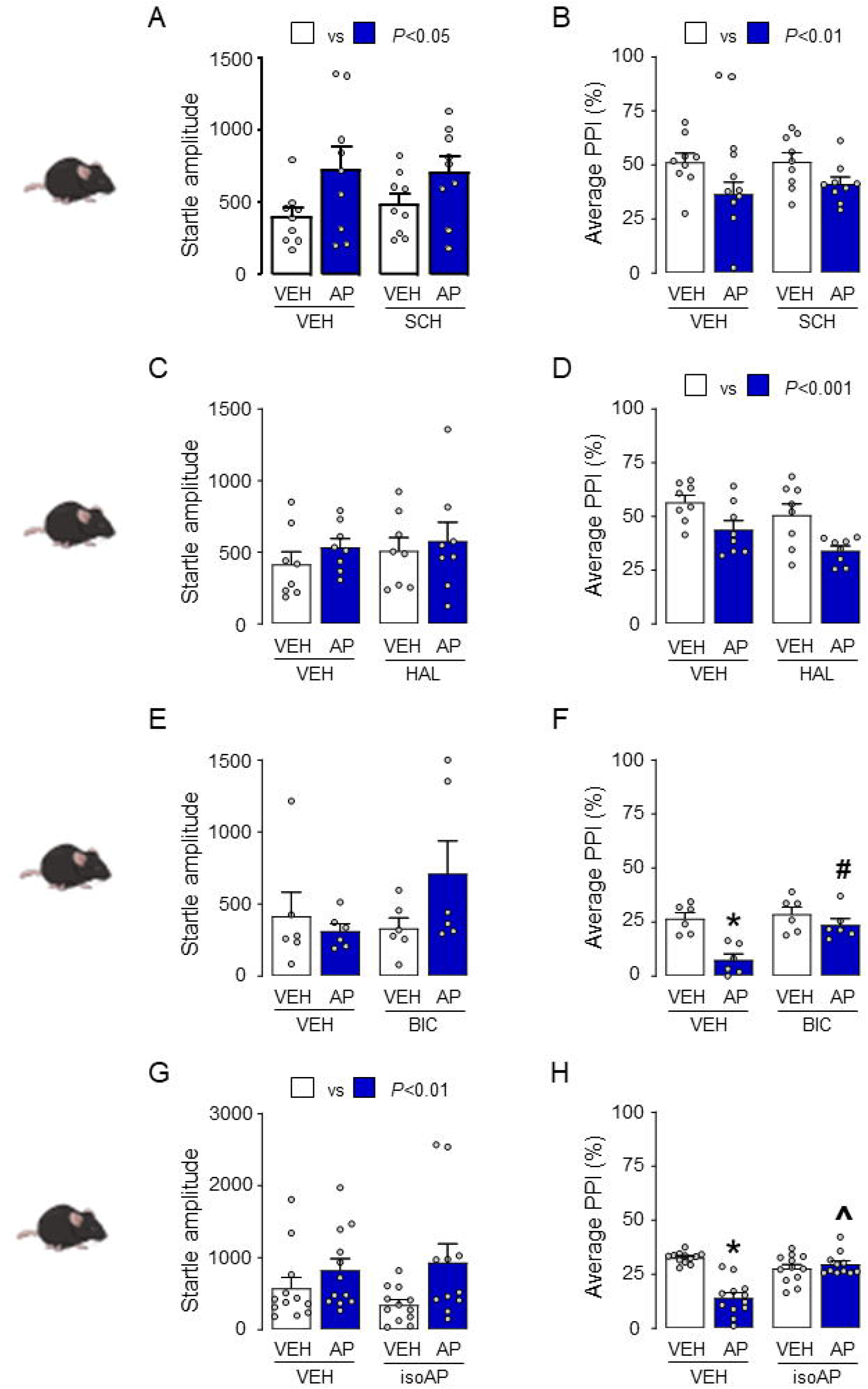
Effects of SCH 23390 (SCH; A-B), haloperidol (HAL; C-D), bicuculline (BIC; E-F), and isoallopregnanolone (isoAP; G-H) on the systemic effects of allopregnanolone (AP) on startle reflex and PPI. Main effects of AP (12 mg·kg^-1^, IP) injections are indicated as comparisons between white and blue bars. Data are shown as means ± SEM; n=6-9/group. VEH, vehicle. *, *p* < 0.05 for comparisons between AP-VEH and VEH-VEH; ^#^, *p* < 0.05 for comparisons between BIC-AP and VEH-AP; **^^^**, *p* < 0.05 for comparisons between isoAP-AP and VEH-AP. For further details, see text.

**Figure 4.**
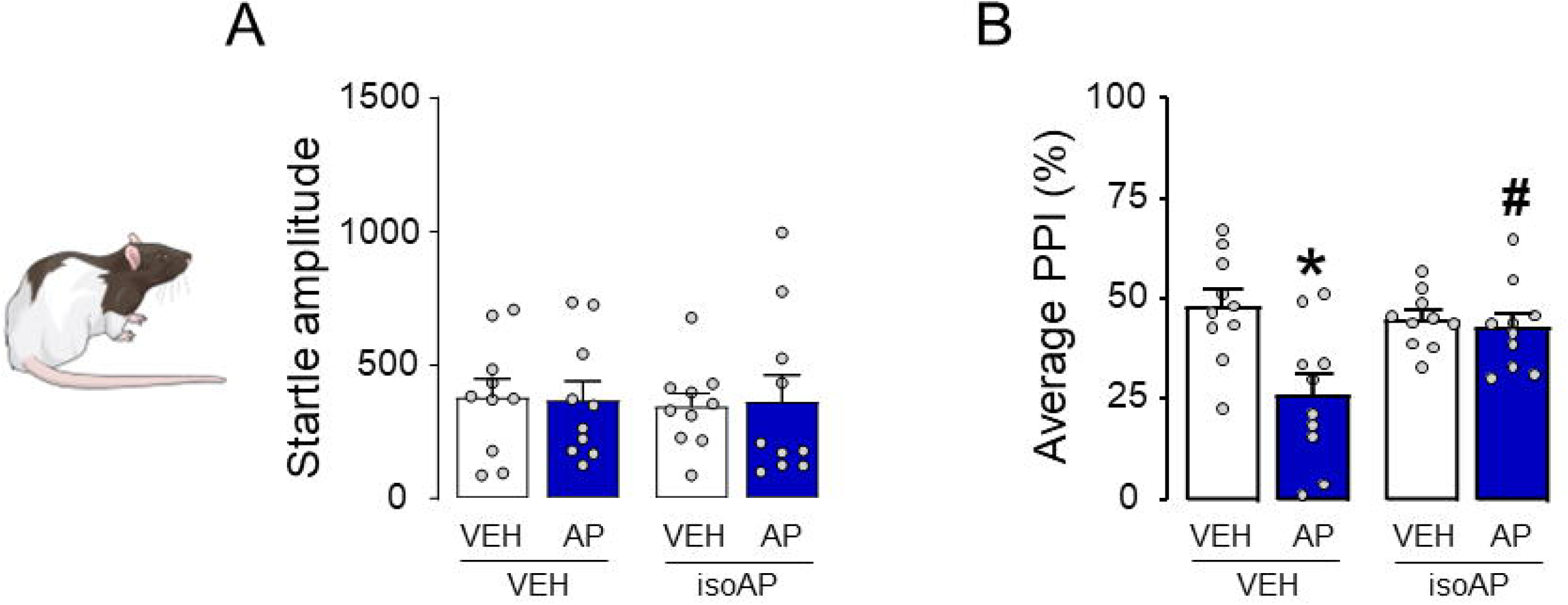
Modifications of startle and prepulse inhibition (PPI) by allopregnanolone (AP; 12 mg·kg^-1^) and its antagonist isoallopregnanolone (isoAP; 10 mg·kg^-1^), administered systemically. Data are shown as means ± SEM; n=10/group. *, *p* < 0.05 compared with rats treated with vehicles (VEH) of AP and isoAP; ^#^, *p* < 0.05 compared with rats treated with AP and VEH of isoAP. For further details, see text.

### PPI is impaired by cortical but not accumbal AP infusions

To ascertain the neuroanatomical basis of AP’s effects on sensorimotor gating, we tested the impact of local infusions of this neurosteroid into the medial prefrontal cortex (mPFC) and nucleus accumbens (NAc), given that these two brain regions were previously shown to contribute to the role of neurosteroids in PPI regulation (Devoto et al., 2012). Bilateral infusions of AP (0.25-1 μg/mouse, delivered as 0.125-0.5 μg/side, within 1 μL of vehicle) in the mPFC did not modify the startle amplitude (Fig. 5A) but reduced PPI (One-way ANOVA: F_3,43_=4.58, *p* < 0.05; *post-hoc* comparisons, vehicle vs. 1 μg AP group, *p* < 0.05; Fig. 5B). Conversely, infusions of AP in the NAc failed to modify either index Figs. 5C-D). To verify whether the mPFC mediates the PPI-disruptive effects of AP, we tested whether intra-PFC injections of isoAP (1 μg/mouse delivered as 0.5 μg/side, 5 min before AP) could counter the effects of systemic AP (12 mg·kg^-1^, IP) (Figs. 5E-F). While no significant modifications of the startle were observed (Fig. 5E), the PPI deficits induced by systemic AP were fully reversed by intra-PFC iso-AP (Two-way ANOVA; AP × isoAP interaction: F_1,28_=18.61, *p* < 0.05; *p*’s < 0.001 for comparisons between vehicle-vehicle and vehicle-AP and between isoAP-AP and vehicle-AP; Fig. 5F).

**Figure 5.**
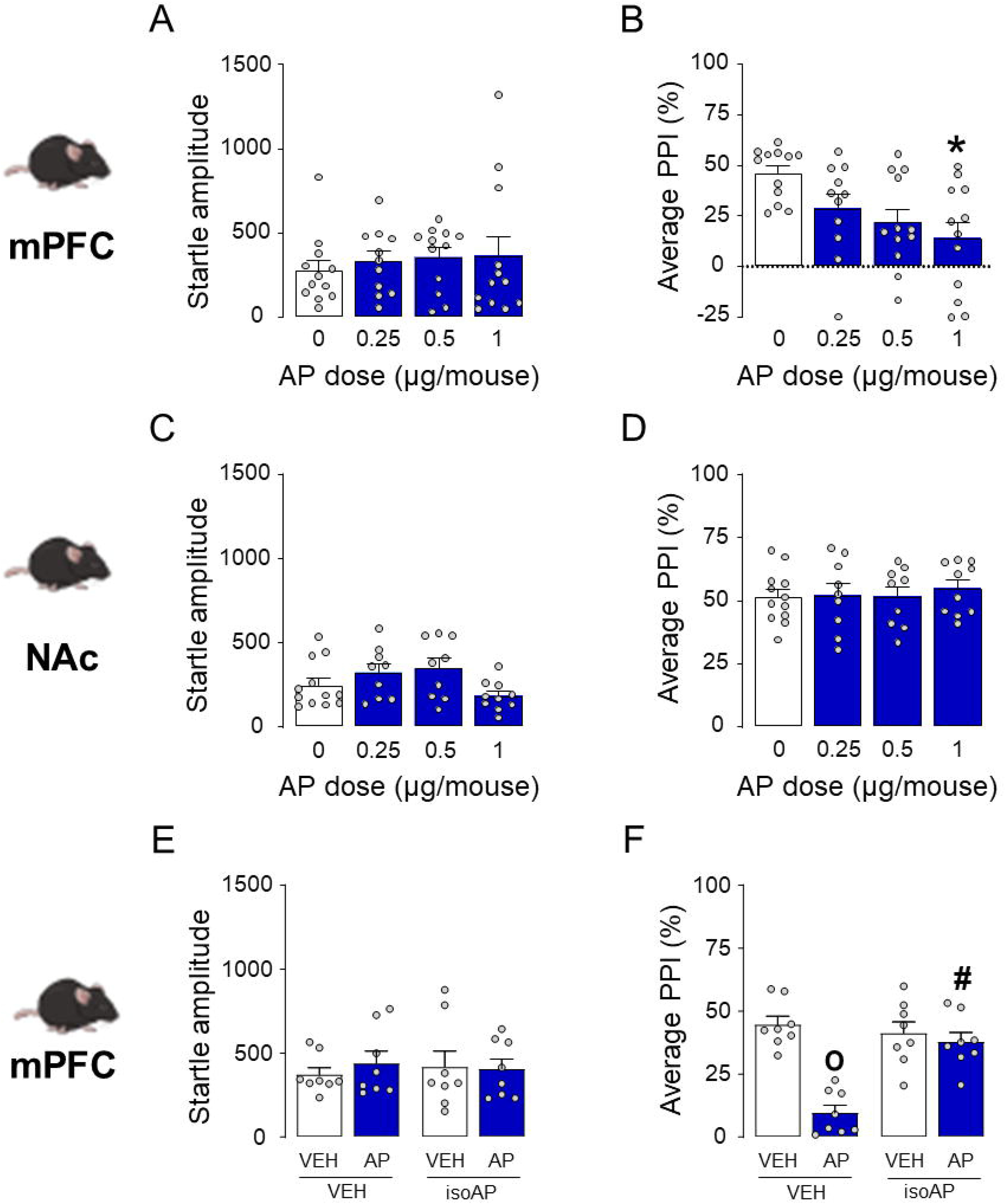
Brain-regional effects of allopregnanolone (AP; 0.25-1 μg/animal) on sensorimotor gating of C57BL/6J mice. AP was infused into the medial prefrontal cortex (mPFC, A-B) and nucleus accumbens (NAc, C-D); then, its antagonist isoallopregnanolone (isoAP) was infused into the mPFC in combination with the systemic effects of AP (12 mg·kg^-1^, IP, E-F). Intracerebral AP doses are indicated below each bar. Data are shown as means ± SEM; n=8-12/group. VEH, vehicle. *, *p* < 0.05 compared with 0 mg·kg^-1^ (VEH-treated); **^°^**, *p* < 0.05 compared with mice treated with VEH of AP and isoAP; ^#^, *p* < 0.05 compared with mice treated with AP and VEH of isoAP. For further details, see text.

### Acute stress impairs PPI in C576BL/6J mice and Long-Evans rats

Since AP plays a crucial role in the orchestration of stress response, we aimed at testing C57BL/6J mice through an array of diverse acute stressors mimicking distinct environmental challenges, namely foot shock, sleep deprivation, restraint, and predator exposure. We first tested the effects of a sequence of nine brief footshocks (intensity: 0.2 or 0.5 mA, duration: 1 s each) in C57BL/6J mice (Figs. 6A-B). This stressor increased startle amplitude (One way ANOVA: F_2,20_= 4.074, *p* < 0.05), although multiple comparisons revealed only a marginal statistical trend for a significant effect of 0.5-mA footshock (*p* = 0.06 between 0.5 and 0 mA; Fig. 6A). In addition, footshock significantly reduced PPI (One way ANOVA: F_2,20_= 7.96, *p* < 0.05) in an intensity-dependent fashion (*p* < 0.05 between 0.5 and 0 mA; Fig. 6B). In contrast with these effects, we found that neither restraint nor predator exposure stress (up to 2 h) significantly modified startle amplitude and PPI (Figs. 6C-F). However, their combination induced a marked PPI deficit (One-way ANOVA: F_3,42_=13.28, *p* < 0.05; *post-hoc* comparisons: rats exposed to stress for 60 and 120 min in comparison to non-stressed rats, *p*’s < 0.05) without affecting startle amplitude (Fig.s 6G-H). We then verified whether C57BL/6J mice might be sensitive to the effects of sleep deprivation since this manipulation was previously found to reduce sensorimotor gating in Sprague-Dawley rats and humans (Frau et al., 2008; Petrovsky et al., 2014). In mice, 24 h of sleep deprivation produced no significant changes in startle amplitude (Fig. 6I) but significantly reduced PPI (Two-way, repeated-measure ANOVA: Time × group interaction: F_1,18_=5.22, *p* < 0.05; *p*’s < 0.05 between sleep-deprived mice and pre-stress baseline or controls after 24-h stress; Fig.6J).

**Figure 6.**
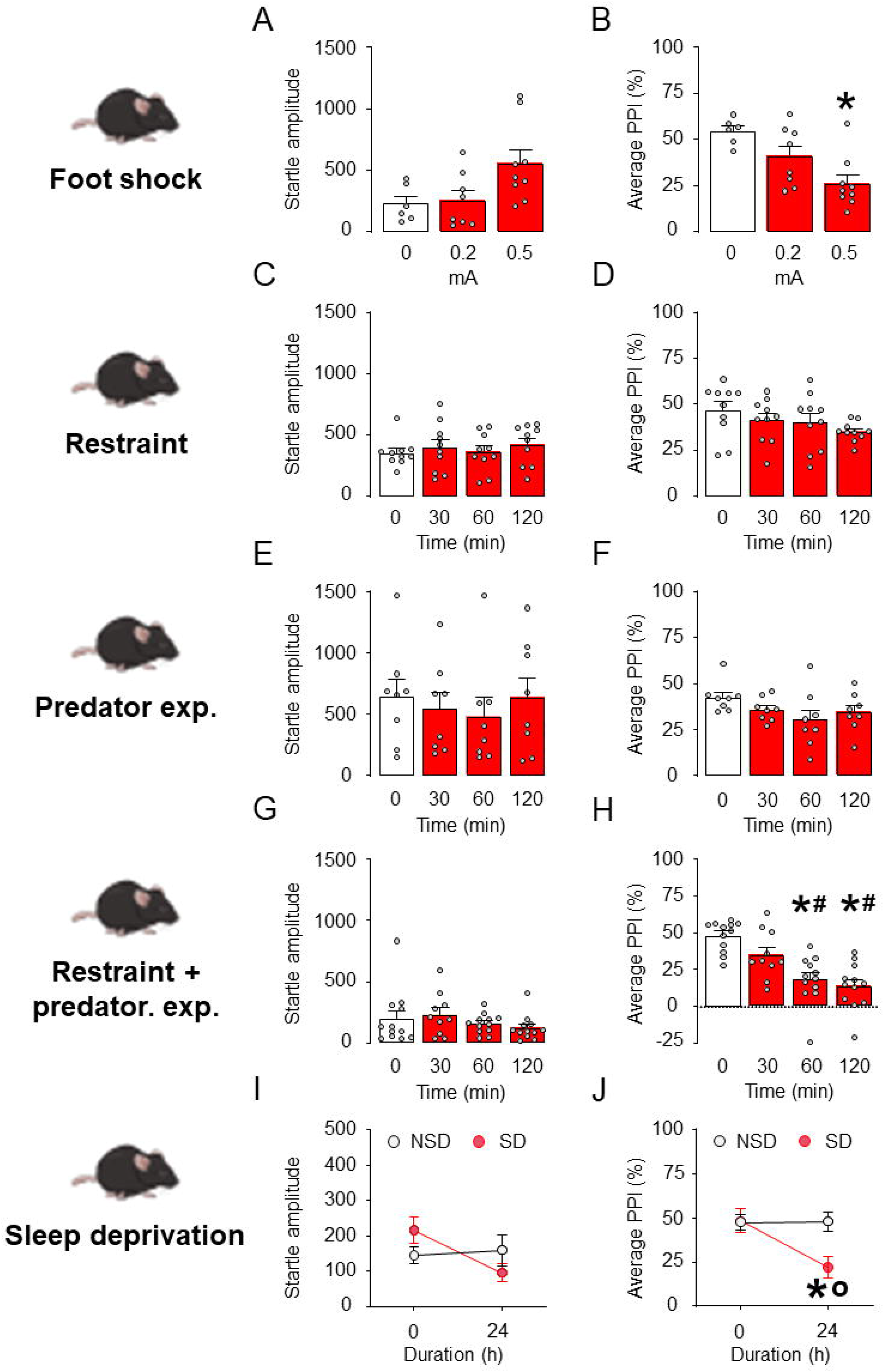
Effect of different stress paradigms on startle and PPI in C57BL/6J mice. A-B) Effects of footshock stress (n=6-8/group). Foot shock intensity is indicated in mA. C-D) Effects of restraint stress (n=10/group). Stress duration of restraint is indicated in min. E-F) Effects of predator exposure stress (n=8/group). Stress duration is indicated in min. G-H) Effects of combined restraint and predator exposure (n=10-12/group). Stress duration is indicated in min. I-J) Effects of sleep deprivation (n=10/group). Stress duration is indicated in h. Data are shown as means ± SEM. *, *p* < 0.05 in comparison with non-stressed rats ^#^, *p* < 0.05 comparison with 30 min restraint; **°**, *p* < 0.05 compared to non-sleep deprived at 24 h. For further details, see text.

Based on these results, we further verified whether the same PPI-disrupting stressors might impair sensorimotor gating in Long-Evans rats. Just as in C57BL/6 mice, a sequence of 0.5 mA foot shocks did not increase startle magnitude (Fig. 7A) but induced PPI deficit (unpaired t-test with Welch’s correction: t= 3.21 df=12.48, *p* < 0.05; Fig.7B). Finally, the combination of restraint and predator exposure failed to modify startle amplitude (Fig. 7C) but produced a time-dependent reduction in PPI (One-way ANOVA: F_2,21_=27.10, *p* < 0.05; *p*’s < 0.05 for *post-hoc* comparisons of 0 and 30 min vs. 60 min,; Fig. 7D). Sleep deprivation did not modify startle amplitude, and this effect did not reflect the duration of this manipulation (Fig. 7E). However, this stressor produced a stable reduction in PPI after 48 h (Two-way, repeated-measure ANOVA: Time × group interaction: F_2,18_ =7.52, *p* < 0.05; *p*’s < 0.05 for *post-hoc* comparisons between sleep-deprived mice and pre-stress baseline or controls after 48-h stress; Fig. 7F).

**Figure 7.**
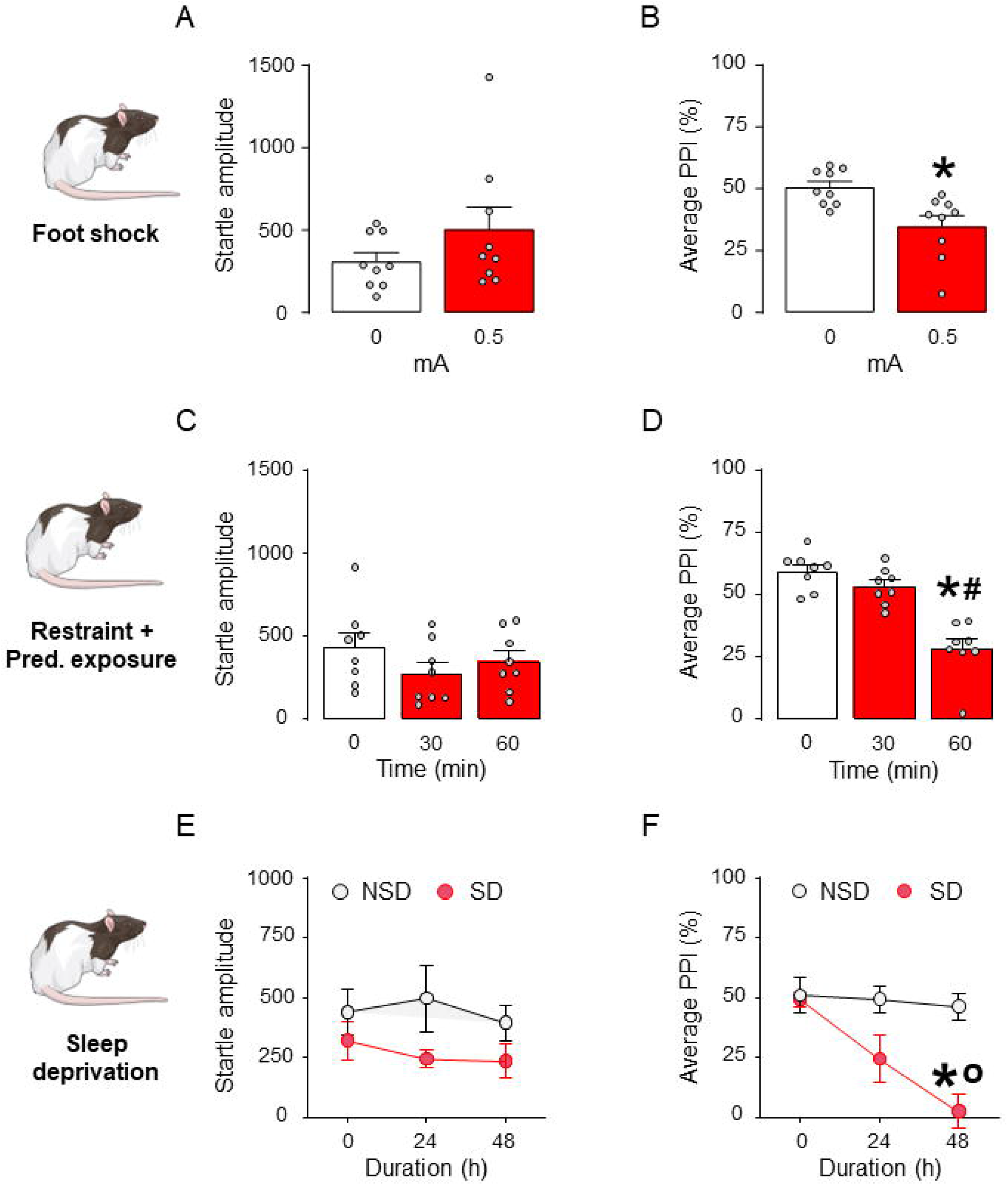
Effect of different stress paradigms on startle and PPI in Long-Evans rats. A-B) Effects of footshock stress (n=9/group). Foot shock intensity is indicated in mA. C-D) Effects of combined restraint and predator exposure (n=8/group). Stress duration is indicated in min. E-F) Effects of sleep deprivation (n=5-6/group). Stress duration is indicated in h. Data are shown as means ± SEM. *, *p* < 0.05 in comparison with non-stressed rats ^#^, *p* < 0.05 in comparison with 30 min restraint; **°**, *p* < 0.05 compared to non-sleep deprived at 24 h. For further details, see text.

### Acute stress increases AP levels in the rat PFC

Building on these results, we tested whether acute stress increased the concentrations of AP (as well as its precursors progesterone and DHP, as well as its metabolite AP sulfate) in the PFC. While foot shock stress did not modify the levels of progesterone (Fig. 8A) and produced a marginal increase in 5α-dihydroprogesterone (DHP) concentrations (Fig. 8B), it significantly enhanced the levels of AP levels (unpaired t-test with Welch’s correction: t=2.45, df=13, *p* < 0.05; Fig. 8C), but not AP sulfate (Fig. 8D).

**Figure 8.**
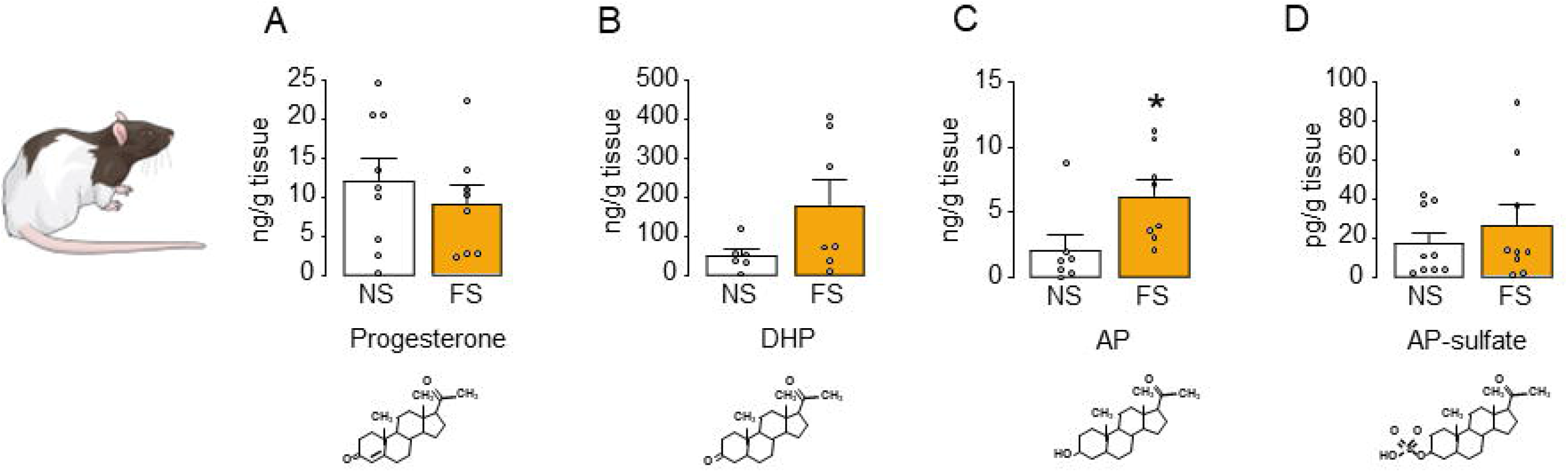
Effects of foot shock (FS; 0.5 mA) on neuroactive steroid levels in the rat prefrontal cortex (PFC). Levels of A) progesterone, B) 5α-dihydroprogesterone (5α-DHP), C) allopregnanolone (AP) and D) AP-sulphate in the prefrontal cortex of Long-Evans rats following footshock stress, compared to non-stressed rats. Data are shown as means ± SEM. n=7-9/group; NS: no stress in comparison to footshock stress, *, *p* < 0.05.

### IsoAP blocks the PPI-disruptive effects of acute stress

The next experiment was aimed at testing whether isoAP was able to counteract the PPI disruption caused by acute stress. In the first set of experiments, the effects of isoAP (10 and 20 mg·kg^-1^, IP, administered immediately after stress) were tested on the behavioral effects of footshock (Figs. 9A-B). While this stressor was confirmed to increase startle amplitude (Two-way ANOVA: Stress × isoAP interaction, F_2,30_=5.07, *p* < 0.05; Fig. 9A) and reduce PPI (Two-way ANOVA: Stress × isoAP interaction: F_2,30_=5.29, *p* < 0.05; Fig.9B), these effects were blocked by isoAP (*p*’s < 0.05 for comparisons between stressed animals treated with mg·kg^-1^ isoAP and its vehicle). Intra-mPFC infusions of isoAP (0.5 μg/side, infused immediately after stress; Figs. 9C-D) produced the same effects on PPI (Two-way ANOVA: Stress × isoAP interaction, F_1,20_=4.68, *p* < 0.05; Fig. 9D), without significantly modifying startle (Two-way ANOVA: Stress × isoAP interaction, F_1, 20_=2.989, *p* = 0.09; Fig. 9C). The effects of the AP antagonist were also observed in relation to the effects of the combination of restraint and predatory exposure stress (Fig. 9E-H). While no overall effects were observed on the startle magnitude (Figs. 9E and 9G), stress-induced PPI deficits were reversed by systemic (10-20 mg·kg^-1^, IP, administered after stress) (Two-way ANOVA: Stress × isoAP interaction, F_2,30_=11.30, *p* < 0.05; *p*’s < 0.05 for comparisons between stressed animals treated with each dose of isoAP and its vehicle; Fig. 9F) and intra-mPFC isoAP (Two-way ANOVA; Stress × isoAP interaction: F_1,32_=12.31, *p* < 0.05; *p* < 0.05 for *post-hoc* comparisons between stressed animals treated with isoAP and its vehicle; Fig. 9H). Conversely, intra-NAc microinjections of isoAP did not alter startle amplitude and failed to oppose the PPI deficits induced by this stressor (Fig.S1).

**Figure 9.**
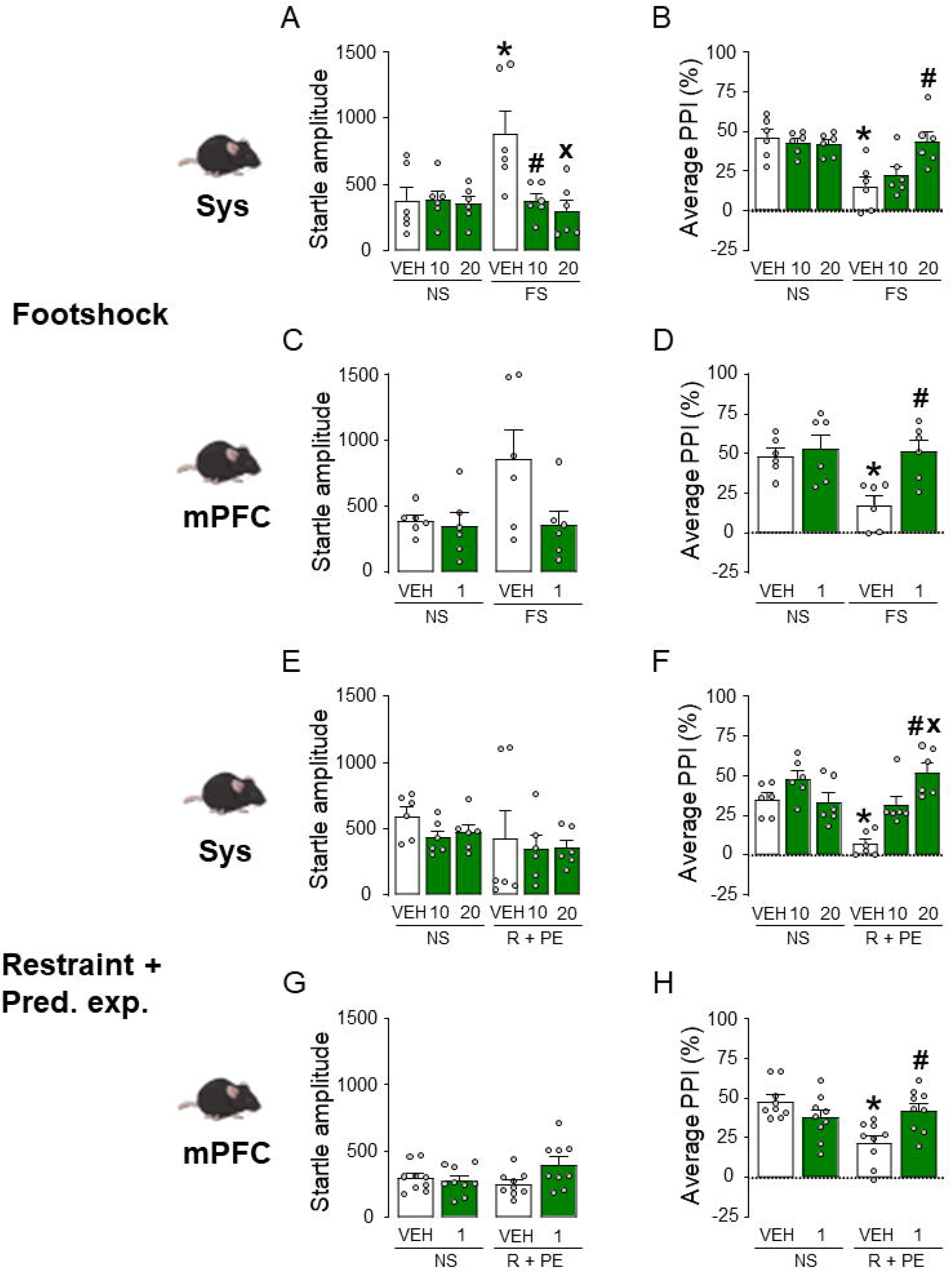
Effects of systemic (SYS) or intra-mPFC isoallopregnanolone (isoAP) on the effects of (A-D) footshock (FS) or (E-H) the combination of restraint (R) and predatory exposure (PE) stress in C57BL/6J mice. IsoAP doses are indicated under the bars, as mg·kg^-1^ (IP) for systemic doses or µg/mouse for local infusions. Data are shown as means ± SEM. n=6-9/group. VEH, vehicle. *, *p* < 0.05 in comparison with VEH-treated, non-stressed (NS) mice; ^#^, *p* < 0.05 in comparison with VEH-treated, stressed mice; **^×^** *p* < 0.05 in comparison with stressed mice treated with isoAP 10 mg·kg^-1^. For further details, see text.

The effects of isoAP were also confirmed in Long-Evans rats (Figs. 10A-D), where this steroid significantly opposed the PPI-disrupting effects of both footshock (Two-way ANOVA; Stress × isoAP interaction: F_2,30_= 3.35, *p* < 0.05 for comparisons between stressed animals treated with 20 mg·kg^-1^ isoAP and its vehicle; Fig. 10B) and 48-h sleep deprivation (Two-way ANOVA; Stress × isoAP interaction: F_1,20_=14.92, *p* < 0.05 for stressed animals treated with 20 mg·kg^-1^ isoAP vs. its vehicle; Fig. 10D) without altering startle amplitude (Fig. 10A and 10C).

**Figure 10.**
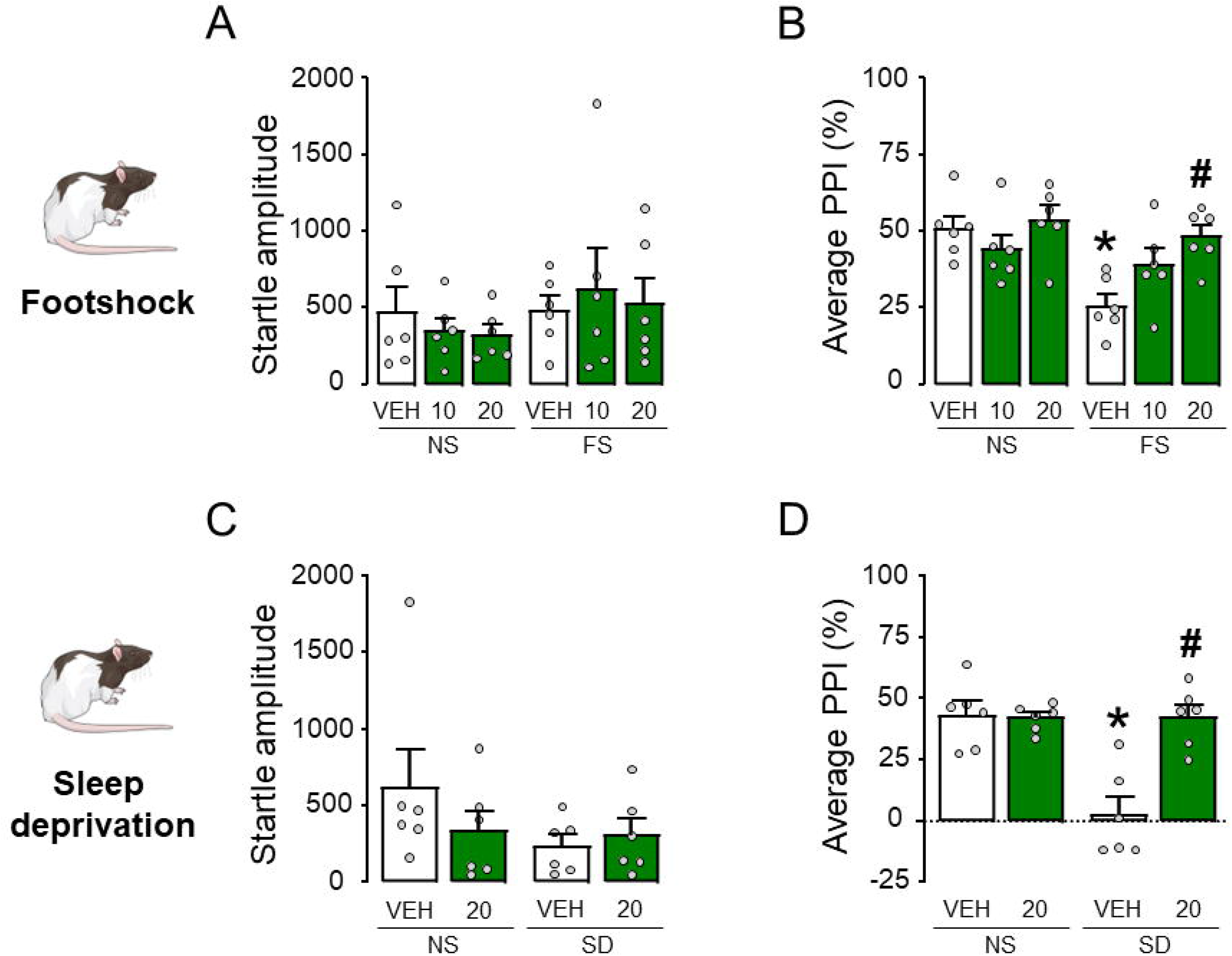
Effects of systemic isoallopregnanolone (isoAP) on the effects of footshock (FS) and 48 h sleep deprivation (SD) stress in Long-Evans rats. IsoAP doses are indicated under the bars as mg·kg^-1^ (IP). Data are shown as means ± SEM. n=6/group. VEH, vehicle *, *p* < 0.05 in comparison with VEH-treated, non-stressed (NS) rats; ^#^, *p* < 0.05 in comparison with VEH-treated, stressed rats. For further details, see text.

## DISCUSSION

The main results of this study showed that systemic and intra-mPFC administration of AP reduced PPI in C57BL/6J mice and Long-Evans rats. Whereas acute stress increased prefrontal AP levels, antagonizing this neurosteroid in the mPFC opposed the detrimental effects of systemic AP injections and stress. Taken together, these results indicate that acute and short-term stress reduces sensorimotor gating by increasing AP in the mPFC. These findings extend previous preclinical evidence suggesting that, although this neurosteroid elicits broad mood-enhancing and anxiolytic properties (Paul and Purdy, 1992; Pinna, 2020), it also exerts a negative influence on cognitive functions, including information encoding (Rabinowitz et al., 2014), spatial learning (Johansson et al., 2002), and memory (Chin et al., 2011; Cushman et al., 2011).

The PPI-disrupting effects of AP were associated with significant or marginal elevations in startle magnitude in a few experiments. These inconsistent effects likely reflect that our protocol was optimized to measure PPI rather than startle magnitude. This issue notwithstanding, the PPI-disrupting effects of AP were not consistently paralleled by changes in startle reflex, indicating that the modifications of sensorimotor gating are not secondary to computational artifacts.

The best-defined mechanism of action of AP is the positive allosteric modulation of GABA_A_receptors (Majewska et al., 1986; Paul and Purdy, 1992; Rupprecht, 2003) through activation of a specific site in the α subunit of these channels (Hosie et al., 2006; 2009). At higher (micromolar) concentrations, AP acts as a GABA_A_ receptor agonist by binding to a second site located at the interface between α and β subunits (Hosie et al., 2009). In line with these mechanisms, the detrimental effects of AP on PPI were opposed by the GABA_A_ receptor blocker bicuculline and isoAP, the selective AP antagonist on this target (Lundgren et al., 2003; Johansson et al., 2016). The finding that bicuculline prevents AP-induced PPI deficits is apparently at odds with previous evidence showing that this drug dose-dependently impairs PPI (Yeomans et al., 2010). However, while the effects of bicuculline reported in this study are likely contributed by the mPFC, the intrinsic effects of this drug on sensorimotor gating reflect the involvement of downstream circuits controlling the execution of the acoustic startle reflex, including the giant neurons in the caudal pontine reticular nucleus (Lingenhohl and Friauf, 1994). Overall, this contrast suggests that GABAergic neurotransmission elicits pleiotropic effects on the modulation of PPI, depending on its distinct actions across different brain regions.

The idea that GABA_A_ receptor activation in the prefrontal cortex impairs sensorimotor gating is in keeping with previous data showing that several information-processing domains are shaped by the contribution of GABAergic signaling in this region (Levar et al., 2019). Given that AP activates both extrasynaptic and synaptic GABA_A_ receptors, and its affinity on different targets is influenced by the specific subunit composition of these ionotropic channels (Belelli and Lambert, 2005), it is likely that sudden elevations of AP in the mPFC may induce distinct alterations of the balance between excitatory and inhibitory neural activity in this region. These changes are predicted to reduce the signal-to-noise ratio in cortical signaling (Dehghani et al., 2016) and interfere with information processing.

Another key finding of our study was that several environmental challenges capturing complementary aspects of short-term stress produced similar PPI deficits. This dose- and time-dependent impairment was observed in both C57BL/6J mice and Long-Evans rats, indicating that a state of high emotional arousal associated with acute, intense, inescapable stress, is likely to attenuate the sensory modulation of startle by interfering with the integrity of perceptual and preattentional filters. In contrast with footshock and sleep deprivation, neither acute restraint alone nor predator exposure (up to 2h) was sufficient to diminish PPI. These results align with previous reports documenting that these two stressors failed to produce an immediate reduction of PPI in rats (Sutherland et al., 2010; Bakshi et al., 2011), even though predator exposure did lead to enduring, albeit delayed, sensorimotor gating deficits (Bakshi et al., 2011). While it could be argued that these results point to possible qualitative distinctions across stress modalities, we also found that the combination of these two manipulations reduced PPI in a time-dependent fashion. Considering that the stressfulness of restraint is arguably increased when combined with fear-induced stimuli (in as far as it prevents any possible attempt to escape), it may be concluded that PPI is disrupted by the intensity of acute stress, rather than its specific source, may be critical to disrupt PPI. This explanation suggests that the negative effects of acute stress on sensorimotor gating may be generalized and depend on the severity of the stress response rather than on the nature of the specific trigger. In line with this interpretation, a large body of human studies has documented that both sensory gating and encoding are similarly weakened by many diverse psychological and physical stressors in humans (Johnson et al., 1993; White and Yee, 1997; Yee and White, 2001; Payne et al., 2007).

Acute footshock stress produced a significant, selective elevation of AP in the PFC. These data agree with previous evidence indicating an increase in brain AP concentrations after footshock (Serra et al., 2002; Pisu et al., 2013), restraint (Higashi et al., 2005), and sleep deprivation (Frau et al., 2017), as well as other acute stressors, such as CO_2_ inhalation (Barbaccia et al., 1996) and forced swim stress (Purdy et al., 1991; Vallée et al., 2000). The augmentation of cortical AP in response to acute stress has been typically interpreted as a physiological reaction to counter its deleterious inflammatory and neurotoxic outcomes (Noorbakhsh et al., 2014; Balan et al., 2021). This framework suggests that PPI deficits may be a trade-off for such beneficial responses. At the same time, this transient information-processing alteration may be construed as advantageous, as it could help prioritize coping resources by readjusting saliency maps. Indeed, previous evidence has shown that these cognitive outcomes are typically associated with the response to acute stress (Hermans et al., 2011; Zhang et al., 2019; Zhang et al., 2022).

Our findings substantiated that both systemic and intra-PFC administration of isoAP effectively opposed the detrimental effects of AP and acute stress on PPI. These results confirm previous findings that underscore the damaging impact of acute stress on sensorimotor gating and show that AP mediates the adverse effects of stress on this information-processing domain. Given the key role of sensorimotor gating in the pathophysiology of several neuropsychiatric disorders, these results agree with previous findings from our group showing that AP mediates the negative effects of stress on tic-like behaviors in models of Tourette syndrome (Mosher et al., 2016) and isoAP reduces these effects (Cadeddu et al., 2020).

In contrast with C57BL/6J mice and Long-Evans rats, the intraperitoneal dose of 10 mg·kg^-1^ AP did not intrinsically diminish PPI in Sprague-Dawley rats, suggesting that this strain has lower sensitivity to the adverse cognitive effects of this neurosteroid. While our data do not elucidate the mechanistic basis of this difference, the present studies revealed that the baseline AP content in the PFC of Long-Evans rats (∼ 2 pg/µg) is about 4-fold higher than its counterpart in Sprague-Dawley rats (Frau et al., 2017). Given that the effects of AP are likely contributed by the sum of its endogenous and exogenous levels, we cannot exclude that administering higher doses of this neurosteroid may also impair PPI in Sprague-Dawley rats. An intriguing corollary of this idea comes from our recent finding that AP is required for the PPI-disrupting properties of D_1_ dopamine receptor activation (Mosher et al., 2019); in line with this concept, D_1_ dopamine receptor agonists reduce PPI in C57BL/6J mice (Ralph-Williams et al., 2003) and Long-Evans, but not Sprague-Dawley rats (Mosher et al., 2016). Future studies are warranted to verify whether heritable variations in AP levels may modulate interspecies and interstrain differences in PPI regulation.

The lower sensitivity of Sprague-Dawley rats to the PPI-disrupting properties of AP may also help explain potential discrepancies between our data and previous reports on the differential effects of stress in this strain. While we documented that footshock engendered PPI deficits in C57BL/6J mice and Long-Evans rats, prior research showed that this stress did not elicit these effects in Sprague-Dawley rats (Bakshi et al., 2011). In line with these ideas, we showed that the duration of sleep deprivation conditions necessary to elicit significant PPI deficits in C57BL6/J mice (24 h) and Long-Evans rats (48 h) was shorter than in Sprague-Dawley rats (72 h; Frau et al., 2008).

Several limitations in the present study should be acknowledged, including our lack of inclusion of female rodents in our studies - which was motivated in view of the significant fluctuations of AP and its precursor progesterone during the ovarian cycle (Genazzani et al., 1995; Palumbo et al., 1995); furthermore, while our data point to a direct action of AP on GABA_A_ receptors, it offers only limited insight into the effects of this process on the molecular signaling and connectivity of the mPFC. Even with these limits, our results support the idea that prefrontal AP mediates the negative impact of acute stress on sensorimotor gating. The fact that this mechanism was validated in two distinct species and strains and in response to multiple acute stressors supports the universal relevance and translational value of these findings. As mentioned above, gating deficits are shared across a broad range of neuropsychiatric conditions, such as schizophrenia, and TS, in which acute stressors are posited to increase symptom severity (Cohen and Docherty, 2004; Ito et al., 2013; Dombrowsky et al., 2014; Godar and Bortolato, 2017). From this perspective, our results appear to point to a crucial role of prefrontal AP in the well-known detrimental effects of stress on the information-processing deficits in these disorders. This interpretation may help explain why finasteride, a neurosteroidogenic blocker that reduces AP synthesis, may elicit therapeutic effects in at least some of these conditions (Koethe et al., 2008; Muroni et al., 2011). Future research will be critical to delineate how changes in AP levels shape the influence of environmental contingencies on the severity of information-processing deficits in neuropsychiatric disorders.

## ACKNOWLEDGEMENTS

This study was supported by the NIH grants R21 NS108722 (to MB)

## AUTHOR CONTRIBUTIONS

MB conceived and designed the study; RC and LJM performed experiments; MB and RC conducted data analysis; MB, RC, and SS wrote the original draft of the manuscript. MB, RC, SS, GMR, PN, and NG reviewed and edited the manuscript.

## DECLARATION OF TRANSPARENCY AND SCIENTIFIC RIGOR

This paper adheres to the principles of transparent reporting and scientific rigor of preclinical research as stated in the *BJP* for Design, Analysis, and Animal Experimentation and as recommended by funding agencies, publishers, and other organizations engaged with supporting research.

---

**Supplementary Figure 1.** Effects of isoallopregnanolone (isoAP) infusions in the nucleus accumbens (NAc) on the effects of the combination of restraint (R) and predatory exposure (PE) stress in C57BL/6J mice. IsoAP dose was 1 μg, delivered bilaterally (0.5 μg/side). Data are shown as means ± SEM. n=8/group. ^&^, *p* < 0.05 for main-effect comparisons between R+PE and non-stressed (NS). VEH, vehicle of isoAP. For further details, see text.

