## Supplementary figures and images for "Acute stress impairs sensorimotor gating via the neurosteroid allopregnanolone in the prefrontal cortex"

### Fig.S1

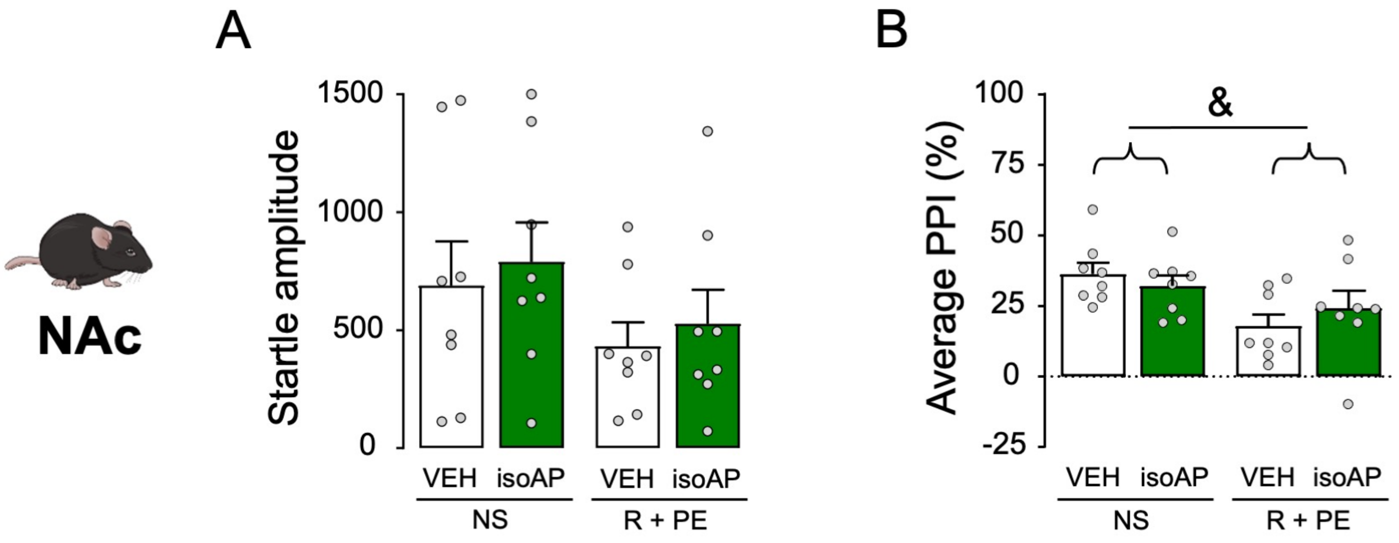
